# Targeted therapy-induced chromosomal instability dictates mitotic dependency on Aurora Kinase A

**DOI:** 10.64898/2026.01.22.701085

**Authors:** Chendi Li, Varuna Nangia, Melissa D. Vieira, Anahita Nimbalkar, Christopher J. Graser, Jeremy Chang, Mohammad U. Syed, Yi Shen, Radhika Koranne, Lee Zou, Xueqian Gong, Sabrina L. Spencer, Aaron N. Hata

## Abstract

Targeted therapies eliminate cancer cells by inhibiting oncogenic signaling; however, tumor cells often evade cytotoxicity through proteomic and epigenetic reprogramming that enables survival. These adaptive responses may create collateral cellular stresses, such as DNA damage, that can be therapeutically exploited. When unresolved, DNA damage leads to chromosomal instability (CIN), a potential source of vulnerability. Whether KRAS inhibition induces DNA damage or CIN in *KRAS*-mutant non-small cell lung cancer (NSCLC) has not been established. Here, we show that the KRAS^G12C^ inhibitor LY3499446 induces CIN in *KRAS*-mutant NSCLC cell lines. A targeted compound screen revealed that the extent of CIN induction by KRAS^G12C^ inhibition strongly correlates with therapeutic synergy with the selective Aurora kinase A inhibitor LSN3321213. Mechanistically, KRAS^G12C^ inhibition stabilizes cyclin B1 during mitosis through activation of mitotic ATR/ATM signaling. In the presence of Aurora Kinase A inhibition, cyclin B1 stabilization delays mitotic exit and diverts cell fate from mitotic slippage or division toward mitotic catastrophe. Together, our findings identify CIN as a predictive marker of response to combined KRAS^G12C^ and Aurora Kinase A inhibition, providing mechanistic rationale to enhance the therapeutic window of AURKA inhibitors when used with targeted therapies.

## Introduction

Most approved targeted therapies eliminate cancer cells by directly inhibiting oncogenic driver proteins. As a consequence of their effects on oncogenic signaling, these agents frequently create collateral cellular stresses, including replication stress^1^, oxidative stress^2,3^, endoplasmic reticulum (ER) stress^4,5^, and various forms of metabolic stress. In particular, recent studies have revealed that targeted therapies such as EGFR tyrosine kinase inhibitors (TKIs)^1,6,7^, BRAF inhibitors^8,9^, or other MAPK pathway inhibitors^10^ can cause DNA damage in cancer cells that survive initial drug exposure by evading apoptosis. These findings highlight that the cellular consequences of targeted therapy extend beyond inhibition of the intended signaling pathway and can involve the induction of diverse stress responses.

KRAS-targeted therapies, particularly KRAS^G12C^ inhibitors (G12Ci), represent a major therapeutic advance but are limited in efficacy. Clinically, G12Ci have shown modest activity in patients with *KRAS*^*G12C*^-mutant non-small cell lung cancer (NSCLC)^11,12,13^, and in preclinical systems such as cell lines and mouse xenograft models, G12Ci often exert primarily cytostatic rather than cytotoxic effects^14-16^. Importantly, previous studies of targeted therapy-induced DNA damage, including with TKIs and MAPK inhibitors, have primarily utilized highly drug-sensitive models, for example, osimertinib treatment in PC9 *EGFR*-mutant NSCLC cells, dabrafenib treatment in DiFi, WiDr, or HT29 colorectal cancer cells, or adagrasib treatment in H358 *KRAS*-mutant NSCLC cells. Far less is known about whether DNA damage can also be induced by targeted therapies in less sensitive or more resistant cancer models, particularly when associated with cytostatic outcomes that more closely mimic clinical responses.

Failure to repair DNA damage prior to mitosis can lead to chromosomal instability (CIN) and chromosomal aberrations^17^. CIN, characterized by an increased likelihood of chromosome mis-segregation during cell division^18^, is considered a hallmark of cancer. CIN has been widely accepted to promote tumor evolution by increasing genomic heterogeneity, enhancing cellular plasticity and adaptability, as well as contributing to therapeutic resistance and metastatic progression^19,20^. At the same time, CIN exposes cells to additional layers of cellular stress, including ER stress, unfolded protein stress, replication stress, proteotoxic stress, and metabolic stress^18^. Because of these vulnerabilities, CIN has been recognized as a potential therapeutic liability, and synthetic-lethal strategies have been developed to target CIN-associated susceptibilities, especially mitotic vulnerabilities^21,22^. However, it remains unknown how KRAS^G12C^ inhibition influences CIN, and whether G12Ci-induced CIN might generate unique therapeutic vulnerabilities that can be exploited.

In this study, we investigated whether G12Ci-induced CIN might be therapeutically exploited in *KRAS*^*G12C*^-mutant NSCLCs representing diverse co-mutation backgrounds. We observed heterogeneous induction of DNA damage and CIN, which positively correlated with enhanced sensitivity to combined inhibition of KRAS^G12C^ and Aurora kinase A (AURKA). Mechanistic studies revealed that G12Ci upregulates mitotic cyclin B1 protein levels through mitotic activation of ATR/ATM DNA repair signaling, thereby prolonging mitotic arrest. Under conditions of AURKA inhibition, in which cyclin B1 degradation is impaired, cells fail to exit mitosis and undergo catastrophic cell death. Thus, our data support a model in which G12Ci-induced CIN creates an Aurora A-dependent mitotic defect that renders *KRAS*-mutant NSCLC cells selectively vulnerable to combined G12Ci + AURKAi therapy.

## Results

### KRAS^G12C^ inhibition induces chromosomal instability and creates a dependency on Aurora kinase A

To determine whether KRAS^G12C^ inhibition can induce DNA damage and CIN, we profiled a panel of 15 NSCLC cell lines harboring *KRAS*^*G12C*^ mutations with diverse co-occurring mutations. Cells were treated with the G12Ci LY3499446 (subsequently referred to as G12Ci), and DNA damage was assessed by γH2AX staining (Fig. 1A). We observed γH2AX-positive cells following G12Ci exposure, with frequencies varying considerably among individual cell lines over time. Some cell lines exhibited sustained increases in γH2AX-positive cells while others displayed only modest or transient increases (Fig. S1A). Since unrepaired DNA damage can lead to chromosome segregation errors and CIN^17^, we also evaluated for the presence of micronuclei (MN), an outcome of CIN (Fig. S1A). Analysis revealed that some cell lines contained Lamin A/C-positive MN (Fig. S1B) at baseline and/or increased numbers of MN after G12Ci treatment, while others did not exhibit significant MN formation (Fig. S1C). Neither γH2AX induction nor MN frequency (baseline or induced) correlated with single-agent G12Ci sensitivity (Fig. S2A-E).

**Figure 1.**
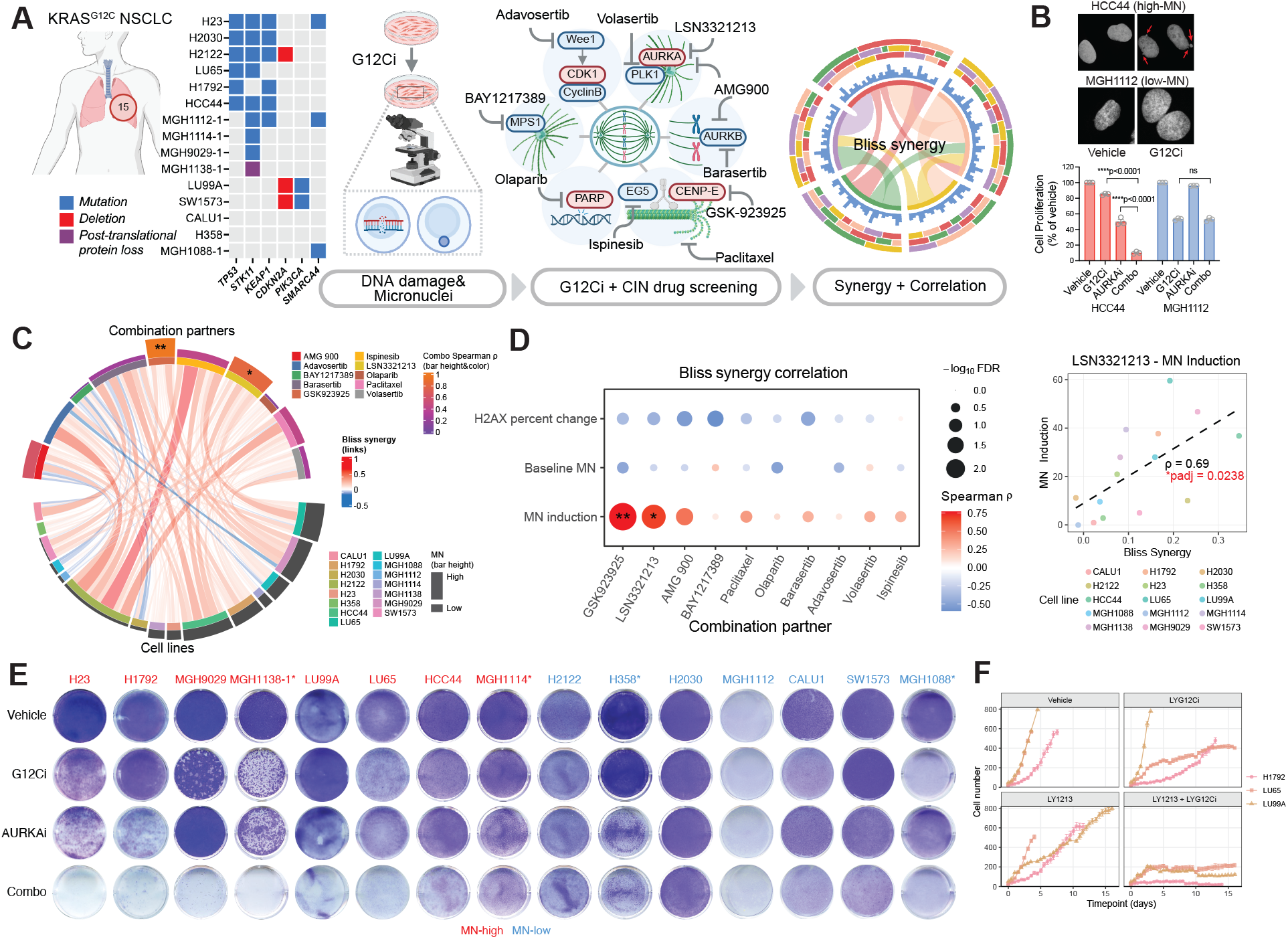
KRAS^G12C^ inhibition induces chromosomal instability and creates a dependency on Aurora kinase A. **A**. Schematic of screening strategy used to identify drug combinations targeting CIN in *KRAS*^*G12C*^-mutant NSCLC cells. **B**. Representative images comparing cell lines with low and high levels of micronuclei (MN) and their corresponding sensitivity to combined LY3499446 (G12Ci) + LSN3321213 (AURKAi). **C**. Chord plot showing synergistic interactions between G12Ci and CIN-targeting drugs for each cell line. The width of each chord represents the Spearman’s rho correlation between synergy and MN formation. Asterisks indicate significant correlations between MN formation and drug synergy. The outer track (upper) denote Bliss synergy and the lower outer track denotes MN induction. **D**. Left: Correlation between Bliss synergy and MN induction, baseline MN, or induction of γH2AX (% change). Asterisks indicate statistically significant correlations. Right: Correlation between MN induction and synergy of LY3499446 with LSN3321213 across 15 *KRAS*^*G12C*^-mutant NSCLC cell lines. **E**. Clonogenic assay showing the effect of 1 µM LY3499446 (G12Ci), 100 nM LSN3321213 (AURKAi) or combination across the cell line panel. Asterisks denote highly sensitive cell lines for which lower G12Ci concentrations (10 nM) were used. MN-high: cumulative MN percent induction > 20. MN-low: cumulative MN percent induction <= 20. **F**. Long-term viability (cell number) of MN-high *KRAS*^*G12C*^-mutant NSCLC cell lines treated with 1 µM G12Ci + 10nM AURKAi.

To investigate whether CIN might confer exploitable vulnerabilities, we assessed drug sensitivity of the 15 cell lines to compounds that target regulators of CIN (Fig. 1A). As expected, drug sensitivity varied across models (Fig. S2F-G). Next, we examined whether targeting these regulators of CIN might exhibit a synthetic lethal interaction with G12Ci. Using sublethal doses for each CIN-targeting agent (specifically determined for each cell line - Fig. S2F-G), we performed drug combination studies with LY3499446. Bliss synergy analysis revealed that many drugs targeting CIN synergized with G12Ci (Fig. S3A). Among these, the Aurora Kinase A inhibitor LSN3321213 (subsequently referred to as AURKAi) exhibited a strong positive correlation between G12Ci synergy and G12Ci-induced MN formation (Fig. 1B-D, S3B). Validation experiments confirmed this relationship: clonogenic assays showed that the G12Ci + AURKAi suppressed colony formation in MN-high cell lines (Fig. 1E, S3C, note that lower G12Ci doses were used for hypersensitive lines), and cytotoxicity assays confirmed that the combination induced cell death rather than merely cytostatic arrest (Fig. S3D). Notably, G12Ci synergized with AURKAi at relatively low nanomolar doses (10nM), leading to substantial suppression of cell viability in MN-high models (Fig. 1F). Together, these results indicate that KRAS^G12C^ inhibition induces CIN in a subset of NSCLC models that confers a selective vulnerability to Aurora kinase A inhibition.

### G12Ci primes Aurora kinase A-dependent mitotic catastrophe

We next sought to define the mechanistic basis for the synthetic lethal interaction between G12Ci and AURKAi. Prior studies have suggested that AURKA can drive adaptive resistance to lung cancer targeted therapies by directly phosphorylating cRAF to sustain MAPK signaling^23^ or phosphorylating and destabilizing the pro-apoptotic protein BIM to suppress mitochondrial apoptosis^21^. When we treated H358 cells with G12Ci, we observed that AURKAi reduced MAPK rebound (p-ERK, p-RSK) after G12Ci treatment, consistent with prior results^23^, however this effect was not consistently observed in other lines that showed sensitivity to the G12Ci + AURKAi combination (Fig. S4A). Additionally, we did not observe increased BIM protein levels in cells upon treatment with G12Ci + AURKAi compared to G12Ci alone (Fig. S4B). We also examined whether prior exposure to G12Ci induced an AURKAi dependent state^23^. We found that when we treated cells with G12Ci for 7 days to select persistent cell populations, they did not become more sensitive to AURKAi or G12Ci + AURKAi compared to treatment-naïve cells (Fig. S4C). Additionally, adagrasib-resistant cells generated by more prolonged treatment with G12Ci did not exhibit increased sensitivity to AURKAi or G12Ci + AURKAi (Fig. S4D), suggesting that sensitivity to G12Ci + AURKAi may not be fully explained by inhibition of adaptive changes.

To gain deeper insight into the synergistic action of G12Ci + AURKAi, we performed lineage tracing using DNA barcoding. We labeled H1792 cells with 100,000 unique DNA barcodes, expanded the population for approximately 20 doublings and divide the barcoded cells into equivalent pools treated with vehicle, G12Ci LY3499446, AURKAi LSN3321213 or the combination. After 14 days of drug treatment, DNA was extracted and barcode sequences were analyzed via PCR amplicon deep sequencing (Fig. 2A). Replicates within each drug condition were highly correlated, indicating non-stochastic clonal selection (Fig. S4E). When examining the top 1000 clones present before treatment, we observed that G12Ci or AURKAi depleted subsets of clones, whereas the combination eliminated the majority of clones that expanded under either single-agent treatment (Fig. 2B). This suggests that AURKAi primes G12Ci-insensitive subclones, or vice versa, to undergo cell death upon dual inhibition.

**Figure 2.**
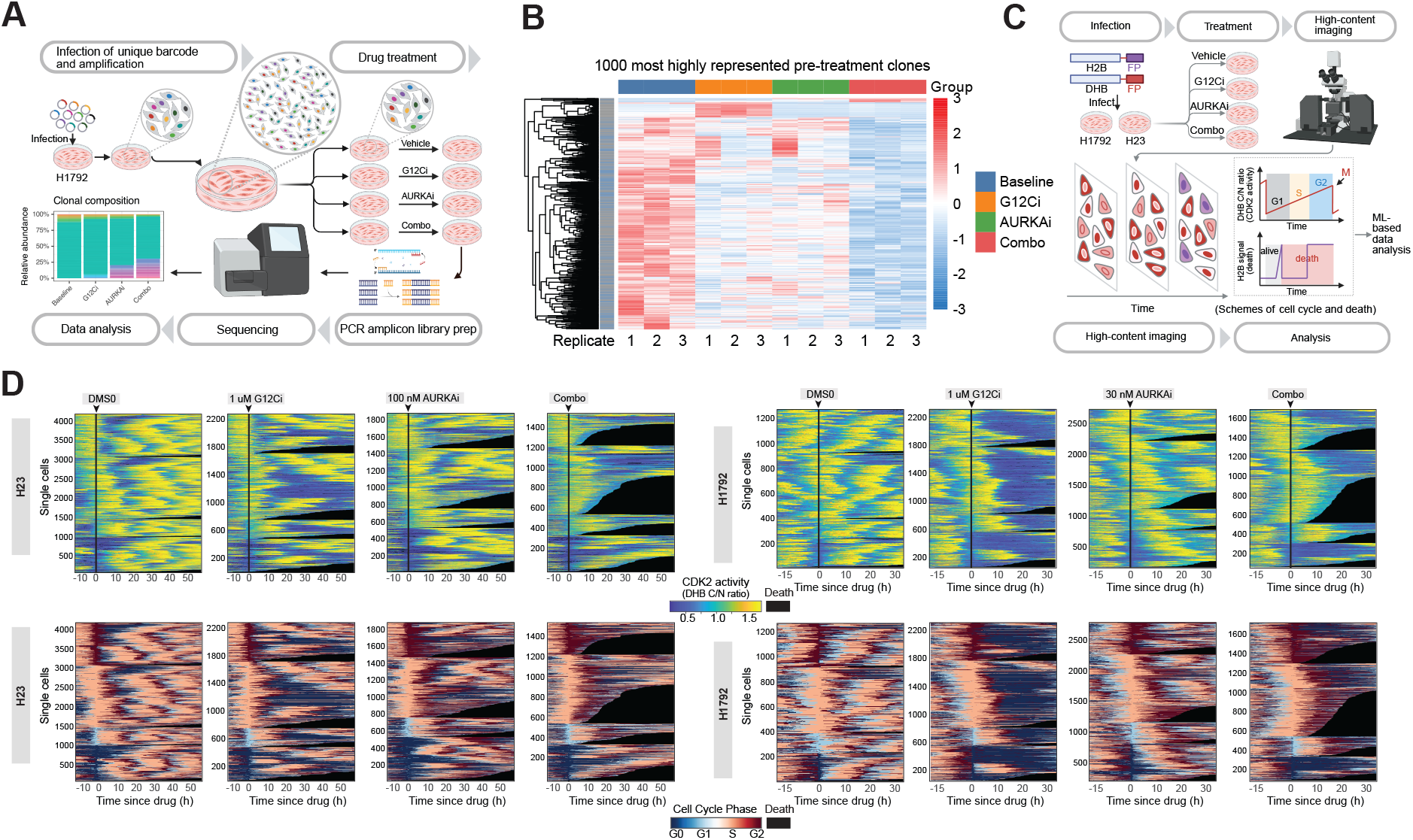
G12Ci primes Aurora kinase A-dependent mitotic catastrophe. **A**. Schematic of ClonTracer barcode lineage tracing experiment. **B**. Top 1000 barcoded clones from H1792 cells treated with vehicle, 1 µM G12Ci, 30 nM AURKAi, or the combination. Higher Z-scores indicate clonal enrichment. **C**. Schematic of single-cell time-lapse imaging using CDK2 and H2B reporters to track cell cycle phase and cell death. **D**. Heatmaps showing single-cell CDK2 activity (upper row) and corresponding cell-cycle phases (lower row) of H23 and H1792 cells treated with vehicle, 1µM G12Ci, 100 nM (H23) / 30 nM (H1792) AURKAi, or combination. Each row represents a single cell. Black tracks indicate cell death. **E**. Representative single-cell CDK2 and H2B reporter tracks showing mitotic catastrophe following G12Ci + AURKAi treatment. **F**. Quantification of cell death across different cell-cycle phases in response to vehicle, G12Ci, AURKAi, or combination treatment.

Given the established role of AURKA in regulating mitotic progression, and our observation of increased combination activity in cell lines with greater G12Ci-induced CIN, we hypothesized that cell proliferation in the presence of G12C might induce an AURKA-dependent mitotic vulnerability. To test this, we performed single-cell time-lapse imaging of H1792 and H23 cells (high synergy between G12Ci and AURKAi), employing a CDK2 activity reporter^24^ to monitor cell-cycle transitions and H2B protein dynamics to track mitotic fate and cell death^10^ (Fig. 2C). Compared to the regular CDK2 oscillations of unperturbed cell-cycle progression in vehicle-treated cells, G12Ci induced G0/G1 arrest and cell death in subset of cells, while some cells continued cycling (Fig. 2D, S4F). AURKAi induced G2/M arrest and partial cell death, while permitting some cells to progress through subsequent cycles (Fig. 2D, S4F). Strikingly, the G12Ci + AURKAi combination triggered a significant increase in cell death, with the majority of death events preceded by cells entering G2/M, as visualized by tracking of characteristic fluctuations between low and high CDK2 activity (Fig. 2D, dark red tracks on lower row heatmaps). Collectively, these results suggest that the synthetic lethality between G12Ci and AURKAi in our models is not driven blocking adaptive MAPK rebound or BIM stabilization, but instead by a G12Ci-induced mitotic defect.

### G12Ci shifts mitotic fate toward catastrophe by stabilizing cyclin B1

We next explored how G12Ci-induced CIN might alter mitotic fate under AURKA inhibition. Prior studies have shown that cancer cells that resume proliferation in the presence of MAPK pathway inhibitors experience replication defects leading to prolonged S phase before mitosis^10^. However, our single-cell tracking experiments did not reveal prolongation of S phase during G12Ci treatment (Fig. S4G), suggesting that G12Ci-induced CIN does not primarily result from replication stress-mediated under-replication of DNA. To determine whether G12Ci treatment altered mitotic progression, we synchronized cells at prometaphase using thymidine block followed by nocodazole treatment (Fig. 3A). Upon release from nocodazole, G12Ci pre-treated cells progressed slightly slower through mitosis with a corresponding decrease in nuclear DNA content compared to vehicle treated cells (Fig. 3B-C, S5B). In contrast, AURKAi pre-treated cells exhibited significantly delayed exit from mitosis and maintained their pre-mitotic nuclear DNA content (Fig. 3B-C). Extended culturing of cells in the presence of AURKAi led to a progressive increase in polyploid cells with up to 8N chromosomal content consistent with mitotic slippage, as determined by flow cytometry (Fig. 3D) and analysis of metaphase chromosome spreads (Fig. 3E, S5A). In contrast, cells treated with the combination of G12Ci and AURKAi underwent mitotic arrest followed by mitotic death (Fig. 3B, S5C). Together, these results reveal that G12C inhibition shifts the mitotic cell fate in AURKAi-treated cells from division or mitotic slippage toward mitotic catastrophe.

**Figure 3.**
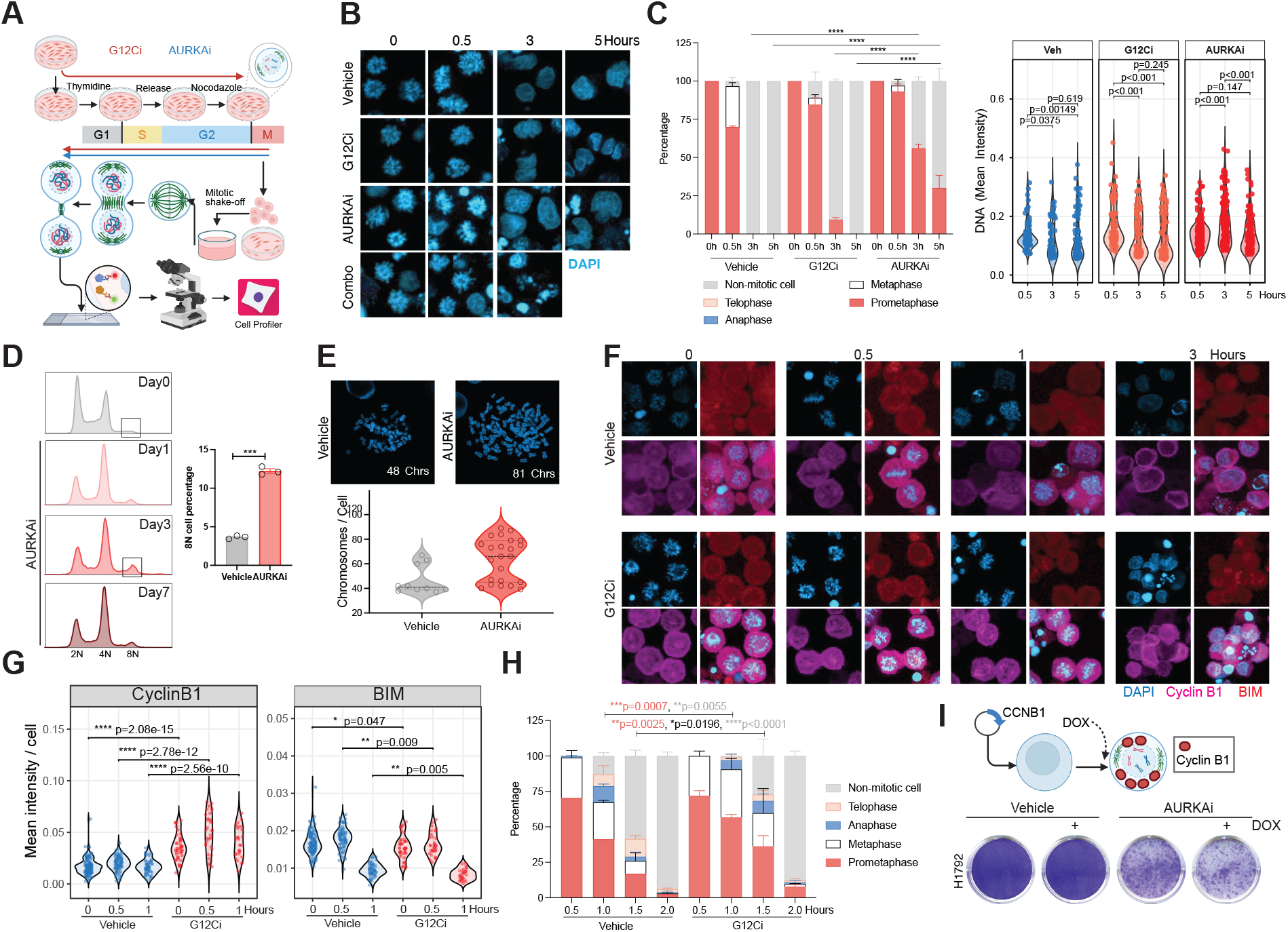
G12Ci shifts mitotic fate toward catastrophe by stabilizing cyclin B1. **A**. Schematic of mitotic synchronization and imaging analysis. **B**. H23 cells were synchronized to prometaphase and released into mitosis. Nuclear morphology was visualized by confocal microscopy. **C**. Top: Quantification of mitotic phase distribution in synchronized H23 cells treated with pretreated with 1 µM G12Ci, 100 nM AURKAi, or combination. Bottom: DNA content of synchronized H23 mitotic cells under the same treatments. **D**. H23 cells cultured in the presence of 100 nM AURKAi for up to 7 and analyzed for ploidy by DAPI staining and flow cytometry. **E**. Top: Representative chromosome spreads of H23 cells cultured in the absence or presence or AURKAi. Bottom: Quantification of 8N cells and chromosome content. **F-G**. Synchronized H23 mitotic cells (pre-treated with vehicle or 1µM G12Ci) were released for the indicated times, and cyclin B1 and BIM protein levels were assessed by immunofluorescence and quantified using CellProfiler. **H**. Quantification of mitotic phase distribution in synchronized H23 cells treated with vehicle or 1 µM G12Ci over a shorter time course. **I**. H1792 and H23 cells with doxycycline (DOX)-induced cyclin B1 expression were treated with 100 nM AURKAi, or AURKAi + 1 µM G12Ci for 14 days, and cell viability was assessed by crystal violet staining.

Mitotic fate is determined by the balance between cyclin B1 degradation and pro-apoptotic signaling: rapid cyclin B1 degradation favors mitotic slippage, whereas sustained cyclin B1 prolongs mitotic arrest and promotes mitotic catastrophe and apoptosis^25^. Consistent with this, mitotic shake-off assays revealed that cyclin B1 was elevated at mitotic entry and remained persistently high in G12Ci-treated compared to vehicle-treated cells (Fig. 3F-G). This stabilization was associated with prolonged early mitosis (Fig. 3H). In contrast, levels of the pro-apoptotic protein BIM were not increased in G12Ci-treated cells during mitosis compared to the vehicle-treated cells (Fig. 3F-G). Overexpression of cyclin B1 in H1792 partially sensitized cells to AURKAi in the absence of G12Ci (Fig. 3I, S5D). (Note: we were unable to over-express cyclin B1 levels in H23 cells). Overexpression of BIM to comparable levels in G12Ci group had minimal sensitizing effect (Fig. S5E-F). These results demonstrate that G12Ci treatment leads to stabilization of cyclin B1, shifting mitotic fate from escape/slippage to catastrophe in the context of AURKA inhibition.

### G12Ci stabilizes Cyclin B1 through ATR/ATM activation in mitosis

DNA damage incurred during mitosis can activate the ATR-CHK1 signaling axis, leading to stabilization of cyclin B1 through modulation of the ubiquitin ligase cofactor Cdh1^26^. To test whether the sensitivity of *KRAS*-mutant NSCLC cells to combined G12Ci + AURKAi is dependent upon DNA damage response signaling, we silenced ATR or ATM in CIN-high cell lines that exhibited strong synergy between G12Ci and AURKAi. Knockdown of either kinase reduced cell death induced by the combination (Fig. 4A, S6A-B), suggesting that sensing of G12Ci-induced DNA damage is critical for altered mitotic fate in response to AURKA inhibition. Interestingly, silencing of downstream checkpoint kinases CHK1 or Wee1 did not similarily blunt the phenotype but instead resulted in profound synthetic lethality (Fig. S6C), consistent with a broader role for CHK1-Wee1 signaling. To test whether DNA damage-induced ATR/ATM signaling can confer AURKA dependency, we treated cells with low-dose doxorubicin to induce MN formation and activate ATR signaling^27^. In ATM-deficient H23 cells, low dose doxorubicin (which has minimal effect on cell viability), induced micronuclei formation (Fig. S6D) and phosphorylation of ATR and CHK1 and increased sensitivity to AURKAi (Fig. 4B), mimicking the effect of G12Ci. These data suggest that activation of ATR and/or ATM primes G12Ci-treated cells for Aurora A-dependent mitotic catastrophe.

**Figure 4.**
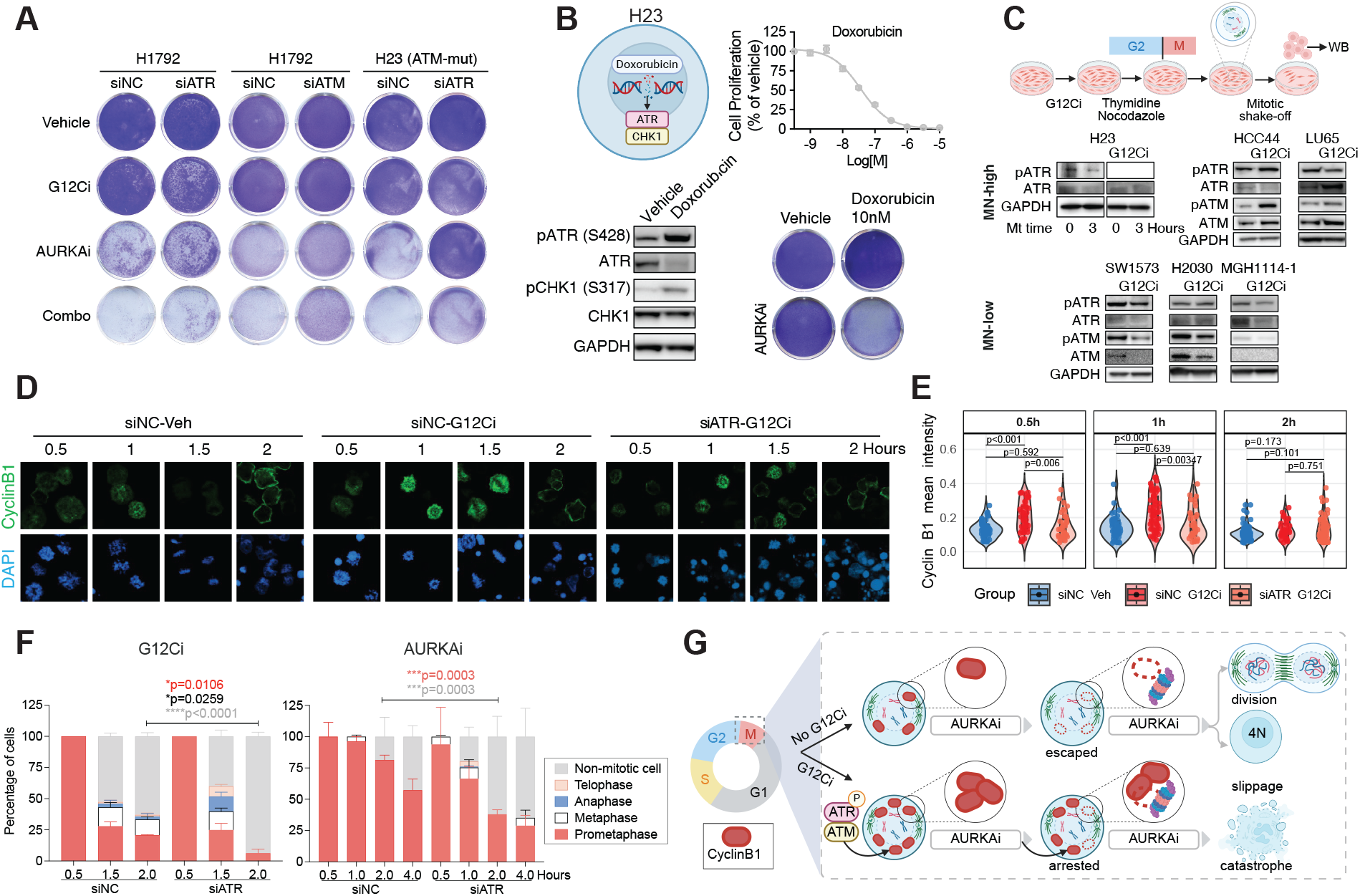
G12Ci stabilizes cyclin B1 through ATR/ATM activation in mitosis. **A**. Viability of H1792 and H23 (ATM-mutant) cells with siRNA knockdown of ATR or ATM (or non-targeting control, siNC) after treatment with G12Ci, AURKAi or combination for 14 days, as determined by crystal violet staining. **B**. Upper panel: Dose response curve of Doxorubicin in H23 cells. Lower panel: MN-high H23 cells were treated with 10 nM doxorubicin for 2 days prior to harvesting for western blotting. H23 cells were treated with doxorubicin, AURKAi or combination for 14 days prior to fixation and staining. **C**. Cell lines with low or high MN levels were treated with 1 µM G12Ci for 3 days, synchronized to prometaphase and released into mitosis. Phosphorylation of DNA damage response proteins was assessed by western blotting. **D-E**. H23 cells with siRNA knockdown of ATR (or non-targeting control, siNC) were treated with G12Ci for 3 days, synchronized to prometaphase and released into mitosis. Cyclin B1 levels were assessed by immunofluorescence (D) and quantified using CellProfiler (E). **F**. H23 cells with NC or ATR knockdown were pre-treated with G12Ci for 3 days, synchronized to prometaphase, and released into mitosis with G12Ci or AURKAi treatment. Mitotic phase distribution was quantified by DAPI fluorescent imaging. **G**. Model illustrating how G12Ci alters mitotic cell fate from division/slippage toward catastrophe and death.

We next examined whether differential activation of ATR activity correlates with the degree of synergy between G12Ci and AURKAi. We did not observe a significant difference in induction of phospho-ATR levels in combination-sensitive compared to insensitive cell lines after G12Ci treatment (Fig. S6E-F), suggesting that ATR activation in unsynchronized cells is insufficient to predict response. However, when we specifically analyzed mitotic populations isolated after synchronization and mitotic shake-off, we observed G12Ci-induced phosphorylation of ATR and ATM in MN-high lines but not in the MN-low lines (Fig, 4C). Finally, we examined cyclin B1 levels in mitotic cells following ATR silencing. Silencing of ATR expression reversed cyclin B1 accumulation in G12Ci-treated cells during early mitosis (Fig. 4D-E), confirming that ATR is required for G12Ci-induced stabilization of Cyclin B1 during mitosis. Furthermore, ATR-silenced cells escaped both G12Ci-induced mitotic slowing and AURKAi-induced mitotic arrest more rapidly compared with control cells (Fig. 4F), suggesting that the reduction in cyclin B1 level upon ATR knockdown facilitates mitotic progression and allows cells to evade mitotic catastrophe.

Collectively, these findings reveal that G12Ci induces activation of ATR/ATM-dependent DNA damage response signaling during mitosis, which in turn leads to stabilization of cyclin B1. Persistent elevation of cyclin B1 prolongs mitosis, further sensitizing cells to AURKAi-mediated mitotic arrest by preventing escape through mitotic slippage and instead committing cells to mitotic catastrophe (Fig 4G).

## Discussion

Targeted therapies induce tumor cell death by suppressing critical oncogenic signaling, but some cancer cells can adapt and survive despite loss of the targeted pathway^28^. In many cases, this can generate collateral stresses that create new therapeutic vulnerabilities. There is growing evidence that targeted therapies that inhibit receptor tyrosine kinase or related downstream signaling pathways can generate DNA damage in cells that survive initial drug treatment^8,9,29^. Here, we demonstrate that KRAS^G12C^ inhibition can induce CIN in *KRAS*^*G12C*^-mutant NSCLC cells, and that this CIN confers a strong dependency on Aurora kinase A.

Aurora kinase A is a serine/threonine kinase essential for centrosome maturation, spindle assembly, and spindle orientation, and also promotes mitotic entry through PLK1 phosphorylation^30^. While most Aurora kinase inhibitors target both Aurora A and B to varying degrees, the LSN3321213 is highly selective for Aurora A, with ∼1,000-fold higher affinity over Aurora B^31^. Prior studies have shown that inhibition of AURKA can cause mitotic slippage^32,33^, consistent with our observations, and that AURKA activity can contribute to resistance to targeted therapies^21,34,35^. However, these earlier studies focused primarily on a role for AURKA in modulating adaptive oncogenic signaling. Our results reveal a previously underappreciated connection between KRAS^G12C^ inhibition, CIN, and Aurora A-dependent mitotic vulnerability.

Mechanistically, we found that AURKAi-induced mitotic slippage can be converted to mitotic catastrophe by G12Ci-induced ATR/ATM activation and cyclin B1 stabilization. ATR and ATM are central mediators of the DNA damage response^36^. Typically, activation of ATR/ATM in G2 in response to DNA damage delays mitotic entry^37^. Previous studies of mitotic DNA damage have often utilized irradiation or microtubule destabilization, which typically induce a high level of DNA damage^38^. In contrast, our data suggest that G12Ci produces subtle DNA lesions that persist into mitosis, where they activate ATR/ATM. This mitotic ATR/ATM activity stabilizes Cyclin B1, which slows mitotic progression. With intact AURKA function, cells ultimately recover and undergo division. When AURKA is inhibited, cells stall in mitosis and ultimately undergo mitotic catastrophe and cell death. While cyclin B1 stabilization provides a compelling mechanistic explanation, additional pathways may also be involved. For example, ATM-CHK2 can stabilize kinetochore-microtubule attachments through Aurora A, promoting lagging chromosomes^38^, and ATR/ATM activation can directly modulate spindle assembly checkpoint function^39^. Whether similar mechanism may also contribute to the effect of combined G12Ci + AURKAi will require further exploration.

An unresolved question is how G12Ci induces the DNA lesions that prime ATR/ATM activation during mitosis. In *KRAS*-mutant pancreatic models, MEK inhibition was shown to disrupt nucleotide biosynthesis and cause replication stress^40,41^. It is possible that specific genetic or epigenetic contexts render certain NSCLCs particularly vulnerable to such nucleotide imbalance, explaining why only a subset exhibited CIN and synergy with AURKAi. Alternatively, these lines may harbor defects in G2/M checkpoints, allowing unrepaired lesions to persist into mitosis. These hypotheses merit future study.

From a broader perspective, it remains to be determined whether CIN induction is a general consequence of targeted therapies or a feature unique to specific oncogenes or disease contexts. Previous work has associated mutant RAS with mitotic stress^42^, and expression of mutant RAS can exacerbate mitotic errors even in otherwise wild-type cells^43^. Other studies have demonstrated that EGFR inhibitors induce chromosome segregation errors in EGFR-mutant lung cancer cells^21^. Thus, targeted-therapy-induced CIN may be context-dependent. Furthermore, while our data highlight Aurora kinase A as a critical liability, other mitotic regulators may also become essential in CIN-high states. While cell cycle-targeting agents may be associated with by dose-limiting toxicities in the clinic^44^, our results suggest that combining such agents with targeted therapies may lower the required doses, expanding the therapeutic window and enabling their clinical use.

Finally, our study highlights the potential importance of stratifying tumors by CIN. We observed that the subset of *KRAS*^*G12C*^-NSCLC models with elevated micronuclei formation exhibited the strongest sensitivity to combined G12Ci + AURKAi. Developing robust clinically-feasible biomarkers to detect CIN in tumors remains a major challenge. Sequencing-based approaches to quantify replication stress and CIN in patient samples are emerging^45,46^, and could provide a path toward guiding patient selection for rational combinations of targeted therapy with agents targeting CIN.

## Supporting information

Supplemental Figure 1 continued

## Acknowledgements

We thank the patients and their families who provided tumor tissue for analysis and generation of patient-derived models. We thank members of the Hata lab and MGH Thoracic Oncology Group for helpful discussion and support. We thank Drs. Stephen Altschuler and Lani Wu at University of California at San Francisco for helpful discussions. This study was funded by support from the NIH F32 CA250231 (C.L.), a research grant from Eli Lilly, and the Ludwig Center at Harvard.

## Author Contributions

C.L., V.N., M.D.V, A.N., M.U.S., Y.S. performed the experiments. C.L., V.N., C.J.G., J.C. performed data analysis and interpretation. X.G. provided KRAS and AURKA inhibitors used in this study. C.L., V.N., S.L.S., A.N.H. were involved with study design. C.L. wrote the manuscript and A.N.H edited the manuscript. All authors discussed the results and commented on the manuscript.

## Methods

### Cell culture

Patient-derived NSCLC cell lines were established in our laboratory from malignant pleural effusions, core-needle biopsies, or surgical resections using previously described methods^47^. All patients provided informed consent under a Dana-Farber/Harvard Cancer Center Institutional Review Board-approved tissue collection protocol and granted permission for research use of their samples. Clinically observed KRAS mutations (determined by the MGH SNaPshot NGS genotyping panel) were verified in the established cell lines. Patient-derived lines were maintained in RPMI supplemented with 10% FBS. Publicly available NSCLC cell lines were obtained from the Center for Molecular Therapeutics at the Massachusetts General Hospital Cancer Center and verified by short tandem repeat (STR) profiling (Biosynthesis, Inc.) at the initiation of the project. Cell lines were routinely tested for mycoplasma contamination during experimental use. Unless otherwise specified, cell lines were cultured in RPMI containing 5% FBS; SW1573 cells were maintained in DMEM/F12 supplemented with 5% FBS.

### Immunofluorescence and image analysis

Cells were fixed with 10% neutral-buffered formalin and permeabilized with PBST (PBS + 0.1% Triton X-100). Cells were incubated with primary antibodies (1:200) overnight at 4 °C, followed by secondary antibody incubation (Thermo Fisher A-11008, A-11012, A-11001; 1:200) for 1 h at room temperature and DAPI counterstaining. Images were acquired on a Zeiss LSM 710 confocal microscope. Image analysis was performed using CellProfiler (Broad Institute). Individual nuclei were identified by DAPI staining, and protein levels were segmented and quantified at the single-cell level. The following primary antibodies were used: Cyclin B1 (Thermo Fisher MA1-155), Lamin A/C (Cell Signaling 4777T), and phospho-histone H2A.X (Ser139) (Cell Signaling 9718S). Downstream analyses were conducted in R.

### Cell proliferation assessment

#### CellTiter-Glo assay

Cell proliferation was measured using the CellTiter-Glo assay (Promega). Cells were seeded into 96-well plates 24 h prior to drug addition. Luminescence was quantified 72 h or 7 days after treatment using a SpectraMax i3x plate reader (Molecular Devices) following incubation with reagent for 30 min at room temperature on a shaking platform.

#### Crystal violet clonogenic assay

Cells in 6-well plates were fixed with 50% glutaraldehyde (Sigma G6403) for 10 min at room temperature, washed twice with water, and stained with 0.1% crystal violet solution (Sigma V5265) for 30 min. After washing and drying, plates were imaged using a flatbed scanner.

#### Incucyte live-cell imaging

Three thousand cells per well were seeded into 96-well plates and treated with indicated drugs. Phase-contrast images were collected using the Incucyte CX3 (Sartorius). Cell numbers were quantified using the cell-by-cell analysis module, and downstream analysis was performed in R.

#### Cytostatic vs. cytotoxic analysis

NSCLC cell lines were treated in a 9 × 9 concentration matrix of G12Ci and AURKAi for 72 h, and cell viability was assessed by CellTiter-Glo. Baseline viability at day 0 was used for normalization, and data were scaled from 0-100-200% relative to baseline.

### Drug Combination and Bliss Synergy Analysis

Combinatorial drug response assays were performed and quantified synergy using the Bliss independence model. KRAS G12C-mutant NSCLC cell lines were seeded in 384-well plates and treated the next day with backbone G12Ci LY3499446 in combination with secondary compound. Cells were treated for 7 days, and viability was measured using CellTiter-Glo 2.0 (Promega) according to the manufacturer’s instructions. The expected combination effect under Bliss independence was calculated as E_Bliss_=A+B−(A×B). A and B are the fractional viabilities of each single agent at a given dose. The Bliss synergy score at each matrix point was then computed as S_Bliss_=O−E_Bliss_. O is the observed fractional viability for the drug combination. The average Bliss synergy per cell line and drug pair was used for downstream analyses, including correlation with phenotypic parameters such as baseline micronuclei frequency, DNA damage (γH2AX percent change), and treatment-induced micronuclei formation. Correlations were quantified using Spearman’s rho, with p values adjusted for multiple comparisons by the Benjamini-Hochberg method.

### Clon-Tracer DNA barcoding analysis

ClonTracer DNA barcoding was performed as previously described^48^. NSCLC cell lines were infected with a ClonTracer barcode library containing ∼100,000 unique barcodes at an MOI < 0.1 to ensure single-barcode integration per cell. Cells were expanded for several doublings to achieve uniform barcode representation, then treated with the indicated drugs for 14 days. Genomic DNA was extracted, and barcode regions were PCR-amplified using ClonTracer-specific primers. Libraries were sequenced on an Illumina MiSeq to a depth of ∼3 million reads per sample. Reads were processed using the original ClonTracer scripts, and barcode counts were normalized to counts per million (CPM) to adjust for sequencing depth. For each barcode, relative abundance was calculated relative to the baseline population. Rank-abundance distributions and cumulative frequency plots were generated, and the top 1000 clones were used to represent dominant populations.

### High-content live cell imaging and cell cycle analysis

#### Stable cell line generation

Low-passage H1792 and H23 NSCLC cells were transduced with H2B-mIFP lentivirus for cell tracking, as well as DHB-mCherry and PIP-mVenus lentiviruses to monitor cell-cycle progression, as previously described^49^. Transduced cells were sorted by fluorescence activated cell sorting (FACS) to establish the cell line.

#### Live-cell time-lapse imaging and single-cell tracking

Cells were seeded on collagen-coated glass-bottom 96-well plates 24 hours before imaging. Movies were acquired on a Nikon Ti-E with a 10×/0.45 NA objective at 15-minute intervals, using phenol red-free growth media. Cells were maintained at 37°C in 5% CO^2^. When drugs were added, imaging was briefly paused to exchange media. Multi-day movies were tracked and processed using EllipTrack^50^. (GitHub: https://github.com/tianchengzhe/EllipTrack).

#### Cell-cycle phase separation with sensor readout

CDK2 activity was quantified as the cytoplasmic/nuclear (C/N) ratio of the DHB signal, as previously described (cite any recent spencer lab paper using live cell tracking here), with the cytoplasmic mean calculated from the top 50th percentile of a pixel ring outside the nuclear mask. PIP-mVenus intensity was extracted from the same nuclear mask and used to report replication licensing state where the PIP signal is high throughout G1, drops sharply upon S-phase entry, and reaccumulates during G2/M. Because the PIP sensor remains high in G0, G1, and G2/M, PIP alone cannot distinguish whether drug-treated cells with high PIP are quiescent (G0) or actively cycling (G1 or G2/M). To resolve these states, we combined PIP intensity with CDK2 activity, which sharply increases as cells commit to the cell cycle and enter G1^24,49^.

Cell-cycle phase classification was performed using fixed thresholds on both reporters: S: PIP < 40 a.u. G0: PIP-high and CDK2 C/N < 0.6. G1: PIP-high and 0.6 ≤ C/N < 1.0. G2/M: PIP-high and C/N ≥ 1.0. To reduce frame-to-frame noise and enforce biologically plausible ordering (e.g., G1 preceding S, S preceding G2/M), we applied simple temporal consistency rules within each single-cell trace. Categorical phase labels (G0, G1, S, G2/M) were converted to integer codes (0-3) for downstream quantitative analysis.

#### Computational determination of apoptosis

To capture apoptotic cells comprehensively, we included both full-length tracks and fragmented tracks typically excluded in standard analyses.

Nuclear H2B intensity traces were log-transformed, smoothed, and rescaled between 0 and 1. Apoptotic events were identified using a “spike-and-plateau” pattern: cells exhibiting a brief intensity surge followed by a sustained plateau of ≥20 consecutive frames above a fixed threshold (0.65) were scored as apoptotic, as previously described^10^. Fragments lacking this characteristic pattern were excluded. Apoptotic fragments were combined with full-length non-apoptotic tracks to generate a unified dataset. All traces were clustered using hierarchical clustering (Euclidean distance, average linkage) and reordered by optimal leaf ordering for heatmap visualization.

#### Relating cell fate to cell-cycle state at drug addition

To link fate to initial cell-cycle state, we defined a short temporal window around the drug-addition frame and assigned each cell a dominant phase based on the modal phase within this window. Cells were grouped by dominant phase (G0, G1, S, G2/M) and sorted within each group by apoptotic timing. Heatmaps of CDK2 ratios and cell-cycle phases were plotted with time on the x-axis and cells ordered by phase and death time on the y-axis, allowing visualization of dynamic signaling and apoptotic progression across the population.

### Mitotic cell synchronization and image analysis

KRAS-mutant NSCLC cells were cultured to ∼70% confluence and treated with thymidine and G12Ci for 24 h, then released into fresh medium for 8 h. Cells were synchronized in prometaphase by treatment with 50 ng/mL nocodazole. Mitotic cells were collected by mechanical shake-off and released into mitosis in suspension culture with or without AURKAi or G12Ci + AURKAi. At indicated time points, cells were collected for immunofluorescence or western blot analysis.

### Chromosome spread

Mitotic cell fractions were prepared as described above. After nocodazole removal, cells were incubated with colcemid for 4 h, pelleted, and treated with 0.075 M KCl for 20 min at 37 °C. Cells were then fixed with methanol : glacial acetic acid (3:1, v/v) and dropped onto pre-cleaned slides. Dried slides were stained with DAPI and imaged using a confocal microscope at 60× magnification.

### Generation of engineered cell lines

Full-length wild-type Cyclin B1 cDNA was subcloned into pInducer20 (a gift from Lee Zou, MGH) via Gateway cloning from EGFP-Cyclin B1 (Addgene #215049). Lentiviral particles were generated by transfecting HEK293 cells with pInducer20 (empty vector) or pInducer20-Cyclin B1 together with VSV-G (Addgene #8454) and Δ8.91 packaging plasmids using Lipofectamine 3000 (Thermo Fisher). KRAS-mutant NSCLC lines were infected and selected with neomycin/G418 to establish stable DOX-inducible cell lines.

### Quantitative RT-PCR analysis

Total RNA was extracted using the Qiagen RNeasy kit, and cDNA was synthesized using the Transcriptor High-Fidelity cDNA Synthesis Kit (Roche) with oligo-dT primers. Quantitative PCR was performed using SYBR™ Select Master Mix (Applied Biosystems) on a LightCycler 480 (Thermo Fisher) with gene-specific primers (Supplementary Table 2). Relative gene expression was calculated using the ΔΔCt method, normalized to *ACTB*.

### Western Blot analysis

Cells were seeded in 6 cm plates and treated when ∼70% confluent. Cells were washed twice with PBS, lysed on ice, and clarified by centrifugation (14,000 rpm, 10 min, 4 °C). Protein concentration was measured using the bicinchoninic acid assay (Thermo Fisher). Equal protein amounts were resolved on NuPAGE 4-12% Bis-Tris gels (Invitrogen) and transferred onto PVDF membranes (Thermo Fisher). Membranes were blocked in 5% milk in TBS-T, incubated with primary antibody (1:1000 in 1% BSA/TBS-T) overnight at 4 °C, then with HRP-conjugated secondary antibody (1:12,500 in 2% milk/TBS-T) for 1 h at room temperature. After three washes in TBS-T, membranes were developed using SuperSignal West Femto substrate (Thermo Fisher) and imaged on a G:Box Chemi-XRQ (Syngene). Primary antibodies included: phospho-ATR (Ser428) CST 2853; ATR CST 13934; phospho-ATM (Ser1981) CST 5883; ATM CST 2873; Aurora A CST 14475; phospho-c-Raf (Ser338) CST 9427; c-Raf CST 9422; phospho-p90RSK (Thr359) CST 8753; RSK1/2/3 CST 9355; phospho-Erk1/2 (Thr202/Tyr204) CST 9101; Erk1/2 CST 9102; Bim CST 2933; Cyclin B1 (Thermo Fisher MA1-155); β-Tubulin CST 2146; and GAPDH CST 5174.

### siRNA-Mediated Gene Knockdown

siRNA transfections were performed using Lipofectamine RNAiMAX (Invitrogen 13778075) per the manufacturer’s protocol. Cells were plated in antibiotic-free medium and transfected at ∼70% confluence in 6-well, 6 cm, or 10 cm dishes. After 48 h, cells were harvested for proliferation assays, immunoprecipitation, or western blotting. The following siRNAs (Invitrogen) were used: negative control (AM4611); siATR (s536, s534); siATM (s1708); siWEE1 (s21); and siCHEK1 (s503).

### Statistics & Reproducibility

All experiments were performed with at least three biological replicates unless stated otherwise. Technical triplicates were included in high-throughput viability and synergy assays. Pairwise comparisons between two groups were performed using two-tailed Student’s t-test for normally distributed data or Wilcoxon rank-sum test for non-parametric data. For multi-group comparisons, one-way or two-way ANOVA followed by Tukey’s or Šidák’s multiple-comparison test was used. For imaging-based quantifications (e.g., micronuclei, γH2AX, or Cyclin B1 intensity), comparisons were performed per biological replicate using mixed-effects models or Wilcoxon tests with Benjamini-Hochberg (BH) adjustment across timepoints. Associations between Bliss synergy and phenotypic parameters (e.g., baseline micronuclei, drug-induced micronuclei, γH2AX fold change) were analyzed using Spearman’s rank correlation coefficient (ρ). The corresponding p values were computed by permutation testing, and false-discovery rate (FDR) correction was applied using the Benjamini-Hochberg method. Linear regression was used to visualize trend lines and confidence intervals in scatterplots. When multiple comparisons were performed, p values were corrected using Benjamini-Hochberg false discovery rate (FDR) control unless otherwise indicated. Significance levels are reported as: * p < 0.05; ** p < 0.01; *** p < 0.001; **** p < 0.0001.

## Supplemental Figure legends

**Figure S1. KRAS**^**G12C**^ **inhibition induces chromosomal instability and creates a dependency on Aurora kinase A. A**. Fifteen *KRAS*^*G12C*^-mutant NSCLC cell lines were treated with 1 µM G12Ci for the indicated days. DNA damage was measured by phospho-H2A.X (S139) staining, and nuclear morphology by DAPI. Quantification of phospho-H2A.X intensity was performed using CellProfiler. Micronuclei (MN) were manually counted; arrows indicate micronuclei. **B**. H23 and H1792 cells treated with 1 µM G12Ci for 7 days were stained with Lamin A/C to visualize micronuclei. **C**. Quantification of micronuclei percentages across cell lines at the indicated time points.

**Figure S2. KRAS**^**G12C**^ **inhibition induces chromosomal instability and creates a dependency on Aurora kinase A. A**. Three-day dose-response curves of G12Ci (LY3499446) in KRAS^*G12C*^-mutant NSCLC cell lines. **B**. Percent viability after 7 days of 1 µM G12Ci treatment relative to vehicle. **C**. Lack of correlation between G12Ci sensitivity and DNA damage. **D**. Baseline MN percentages across cell lines ranked from low to high. **E**. Lack of correlation between G12Ci sensitivity and either baseline MN percentage or MN induction by G12Ci. **F-G**. Area under the curve (AUC) of dose-response curves for chromosomal instability-targeting drugs across 15 KRAS^*G12C*^-mutant NSCLC cell lines.

**Figure S3. KRAS**^**G12C**^ **inhibition induces chromosomal instability and creates a dependency on Aurora kinase A. A**. Heatmap of Bliss synergy scores for G12Ci in combination with CIN-targeting drugs across 15 KRAS^*G12C*^-mutant NSCLC cell lines. **B**. Correlations between γH2AX induction, baseline MN percentage, or MN induction by G12Ci versus Bliss synergy. Asterisks denote adjusted p-value significance. **C**. Clonogenic assays of four highly G12Ci-sensitive cell lines. 1 µM G12Ci was used in this assay. **D**. Cell lines were treated with increasing doses of G12Ci and AURKAi, and cytostatic versus cytotoxic effects were calculated relative to day 0 viability.

**Figure S4. G12Ci primes Aurora kinase A-dependent mitotic catastrophe. A**. Left: Experimental schematic showing G12Ci treatment with 24-hour AURKAi spike-in. Right: MAPK signaling in KRAS^*G12C*^-mutant NSCLC cell lines in response to 1 µM G12Ci and 100 nM AURKAi, assessed by western blotting. **B**. Cell lines were treated for 2 days with 1 µM G12Ci and/or 100 nM AURKAi, and BIM protein levels were analyzed by western blotting. **C**. H23 and H1792 cells were treated with 1 µM G12Ci alone or in combination with 100 nM AURKAi; G12Ci-treated cells were switched to AURKAi or combination treatment. Cell numbers were quantified by live-cell imaging. **D**. MGH1112, LU65, and MGH1062 cells treated with adagrasib for 6 months were assessed for viability to G12Ci (LY3499446) and AURKAi by CellTiter-Glo. **E**. ClonTracer library similarity was calculated using Bray-Curtis distance. **F**. Top: Representative single-cell H2B reporter tracks showing apoptotic versus non-apoptotic cells. Bottom: Representative CDK2 reporter tracks with thresholds for categorizing cell-cycle phases. **G**. Duration of each cell-cycle phase in H23 and H1792 cells under the indicated treatments.

**Figure S5. G12Ci shifts mitotic fate toward catastrophe by stabilizing cyclin B1. A**. Representative chromosome spreads of H23 cells treated with 100 nM AURKAi for 3 days. **B**. Representative images of different phases of mitotic phases. **C**. Quantification of cell death in mitotic fraction of H23 cells in response to 1 µM G12Ci, 100 nM AURKAi or the combination. **D**. H23 and H1792 cells expressing a DOX-inducible cyclin B1 construct were treated with the indicated concentrations of doxycyclin; cyclin B1 levels assessed by western blotting. **E**. H23 cells expressing a DOX-inducible BIM constructed were treated with the indicated concentrations of doxycyclin; BIM expression was assessed by western blotting. DOX concentrations inducing BIM levels comparable to G12Ci treatment were selected. **F**. Clonogenic assays of H23 cells with DOX-inducible BIM were treated with the indicated drugs for 7 days.

**Figure S6. G12Ci stabilizes cyclin B1 through ATR/ATM activation in mitosis. A**. H1792 and H23 (ATM-mutant) cells with ATM and/or ATR knockdown were analyzed by qPCR for gene expression. **B**. Top: qPCR validation of siRNA-mediated ATR knockdown (siATR #2) in H23 cells. Bottom: Viability of H23 cells with ATR knockdown in response to 1 µM G12Ci, 100 nM AURKAi and combination, measured by crystal violet. **C**. Top: qPCR validation of Wee1 and CHK1 knockdown in H23 and H1792 cells. Bottom: Viability of Wee1/CHK1 knockdown cells under the indicated treatments, assessed by crystal violet. **D**. MN formation in H23 cells treated with low-dose doxorubicin; arrows indicate micronuclei. **E**. Left: Phospho-ATR levels in response to G12Ci and AURKAi assessed by western blotting. Right: Quantification of phospho-ATR normalized to total ATR.

