## Supplementary figures and images for "Targeted therapy-induced chromosomal instability dictates mitotic dependency on Aurora Kinase A"

### Supplemental Figure 1

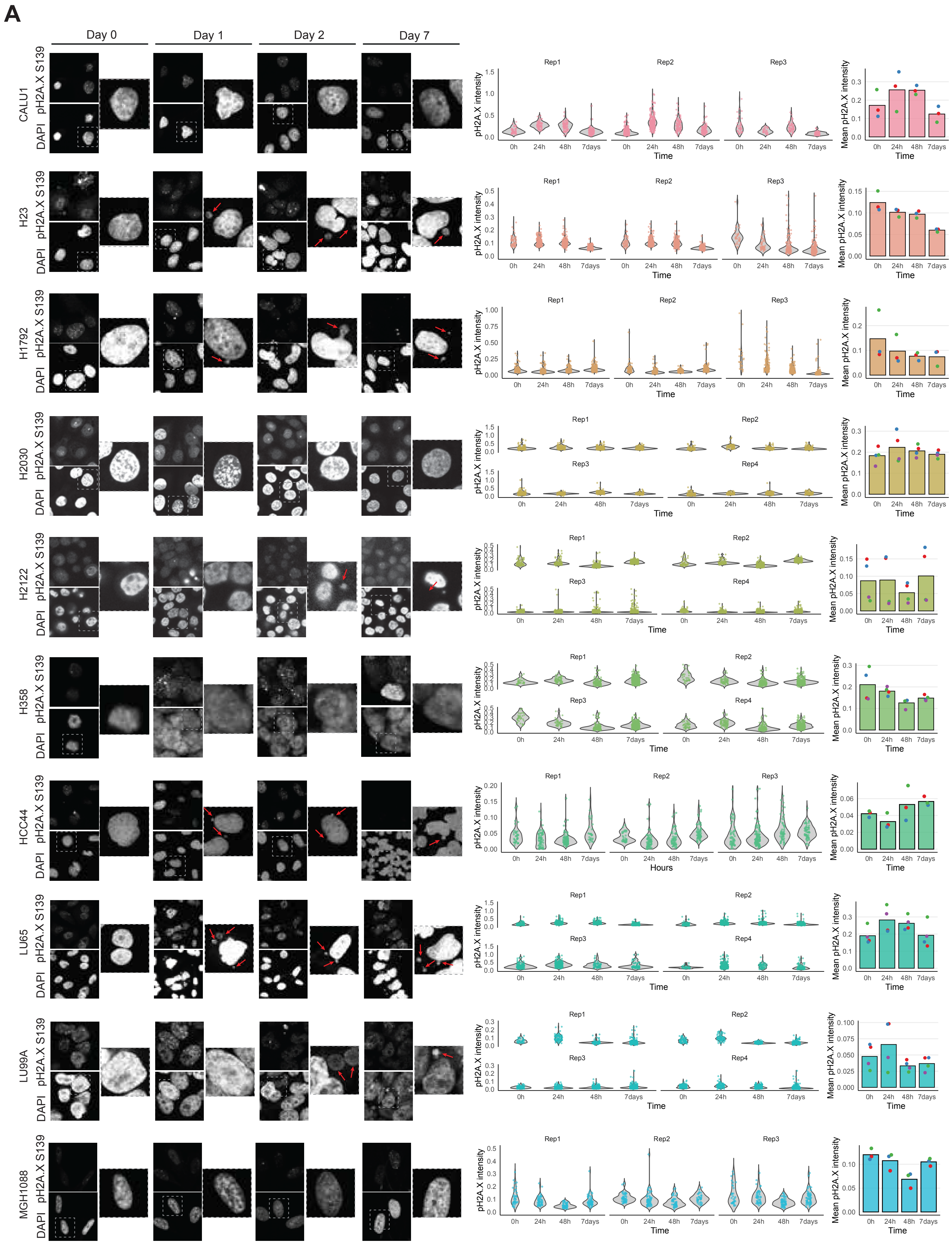


Supplemental Figure 1

### Supplemental Figure 1 continued

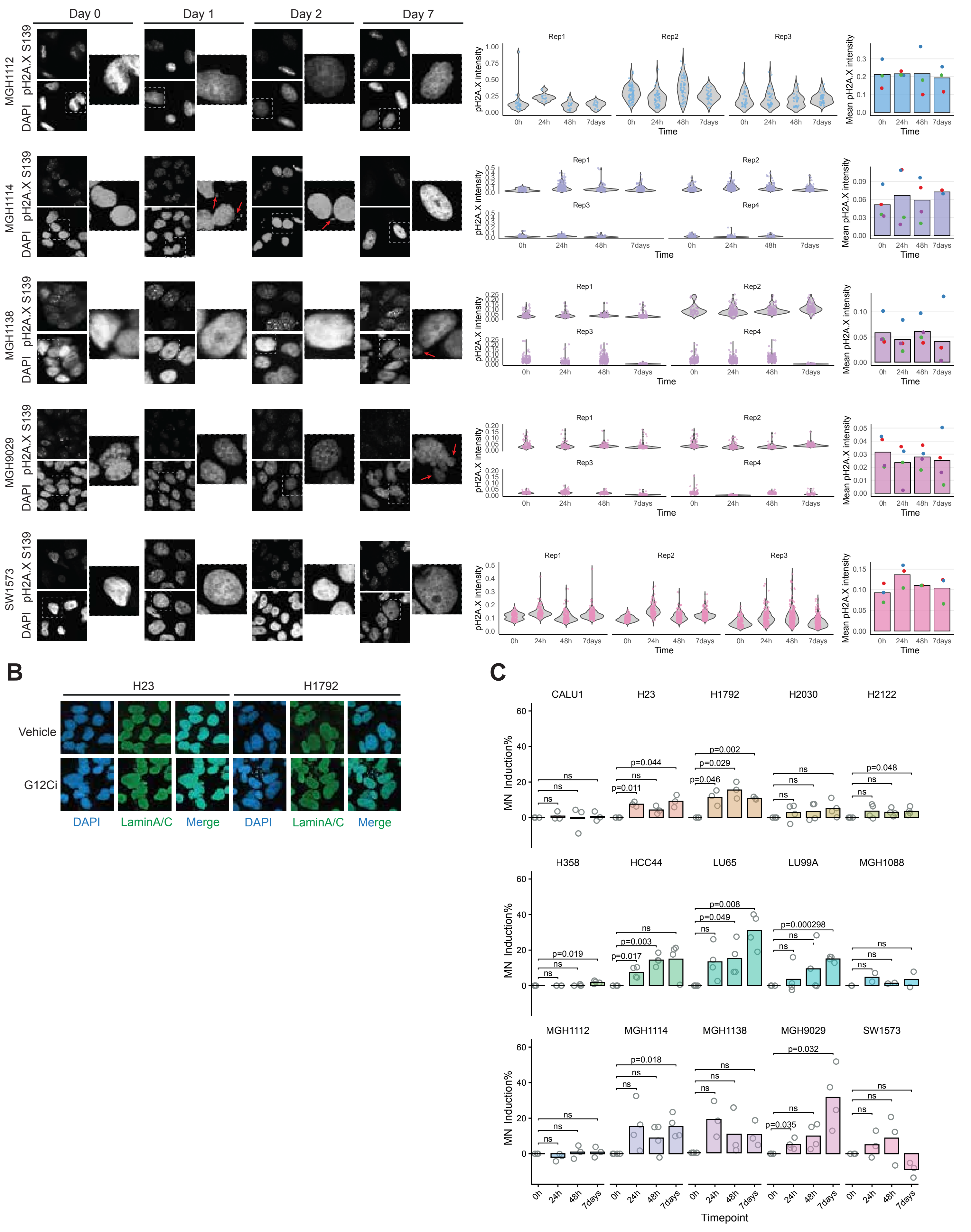


Supplemental Figure 1 continued

### Supplemental Figure 2

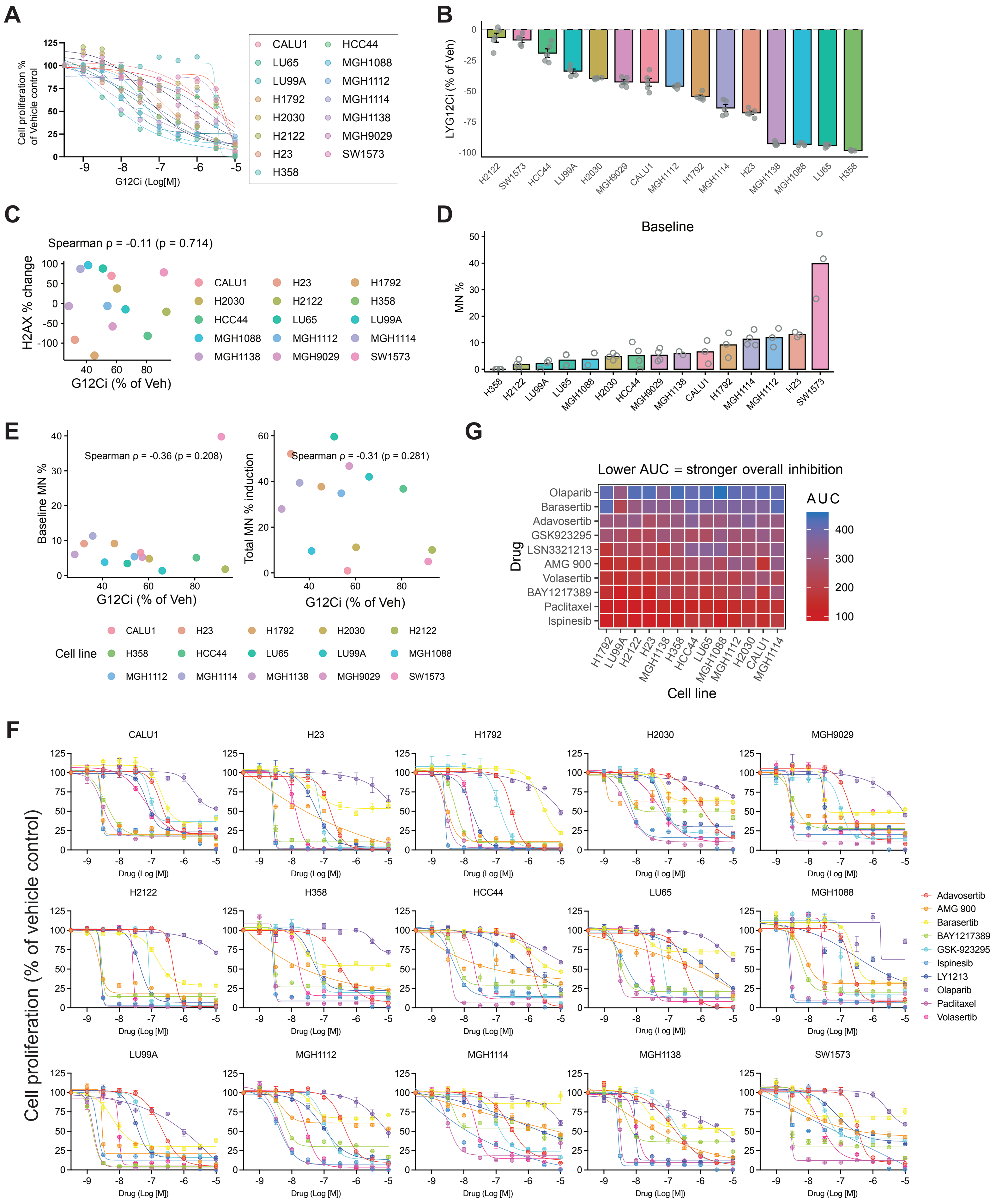


Supplemental Figure 2

### Supplemental Figure 3

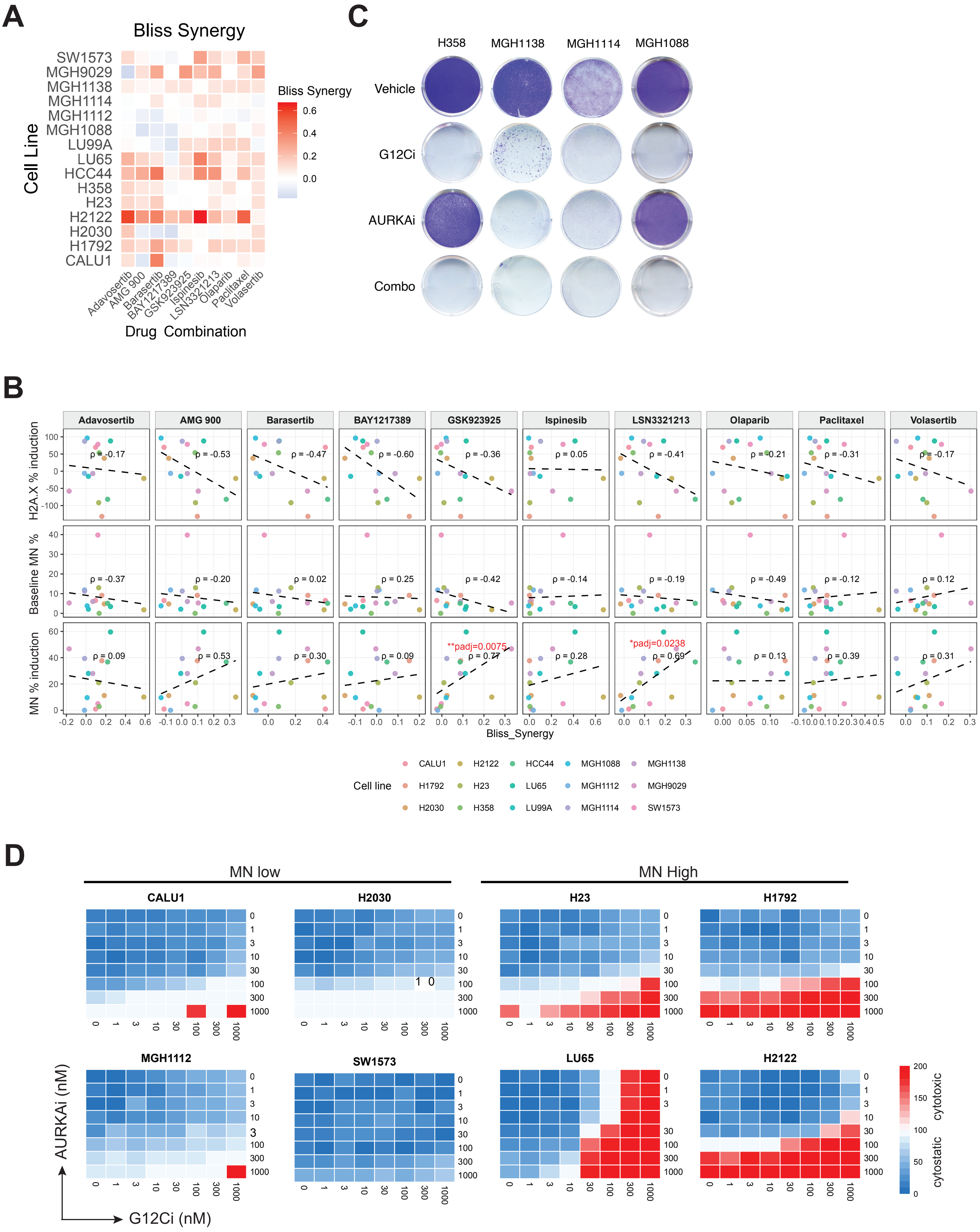


Supplemental Figure 3

### Supplemental Figure 4

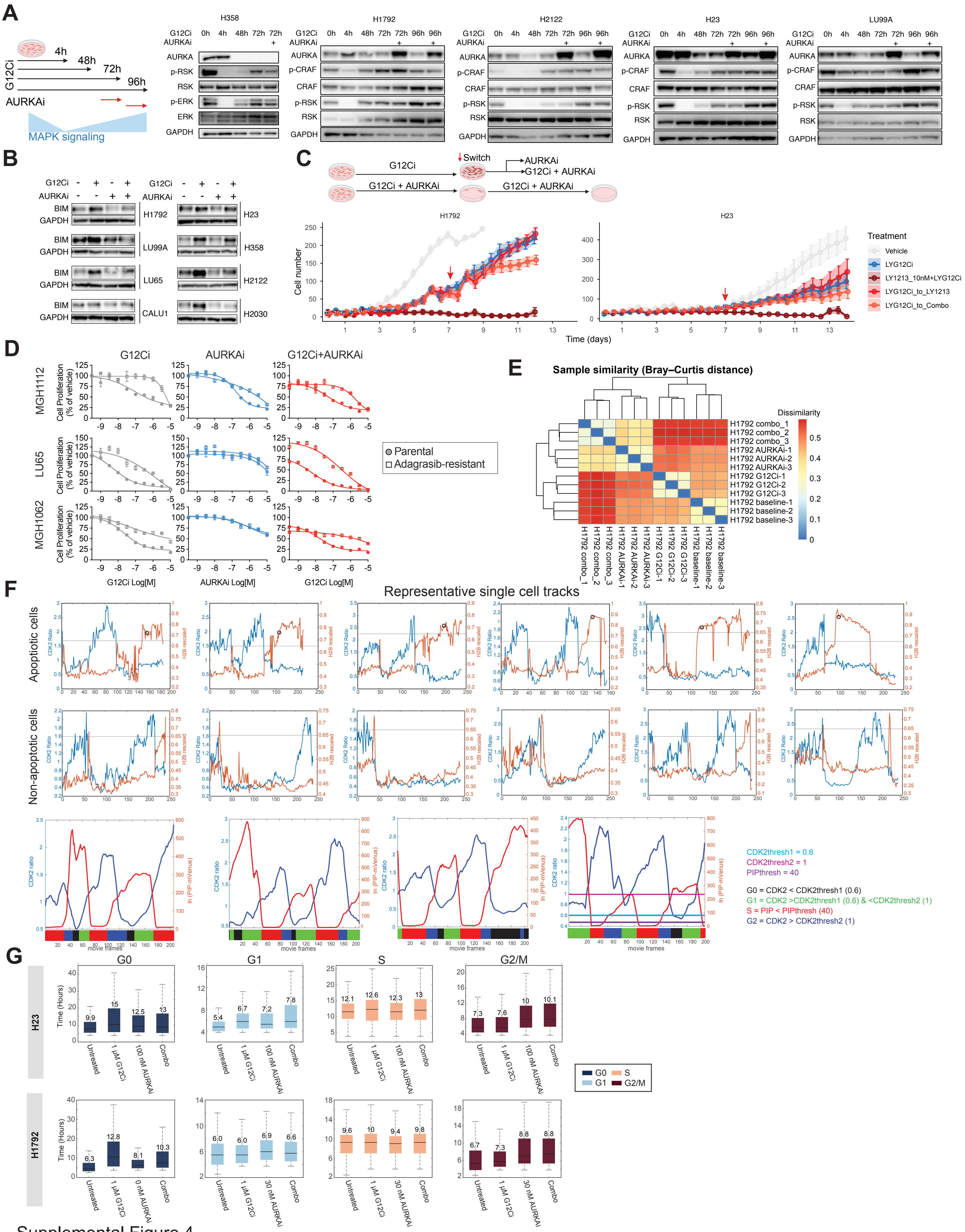


Supplemental Figure 4

### Supplemental Figure 5

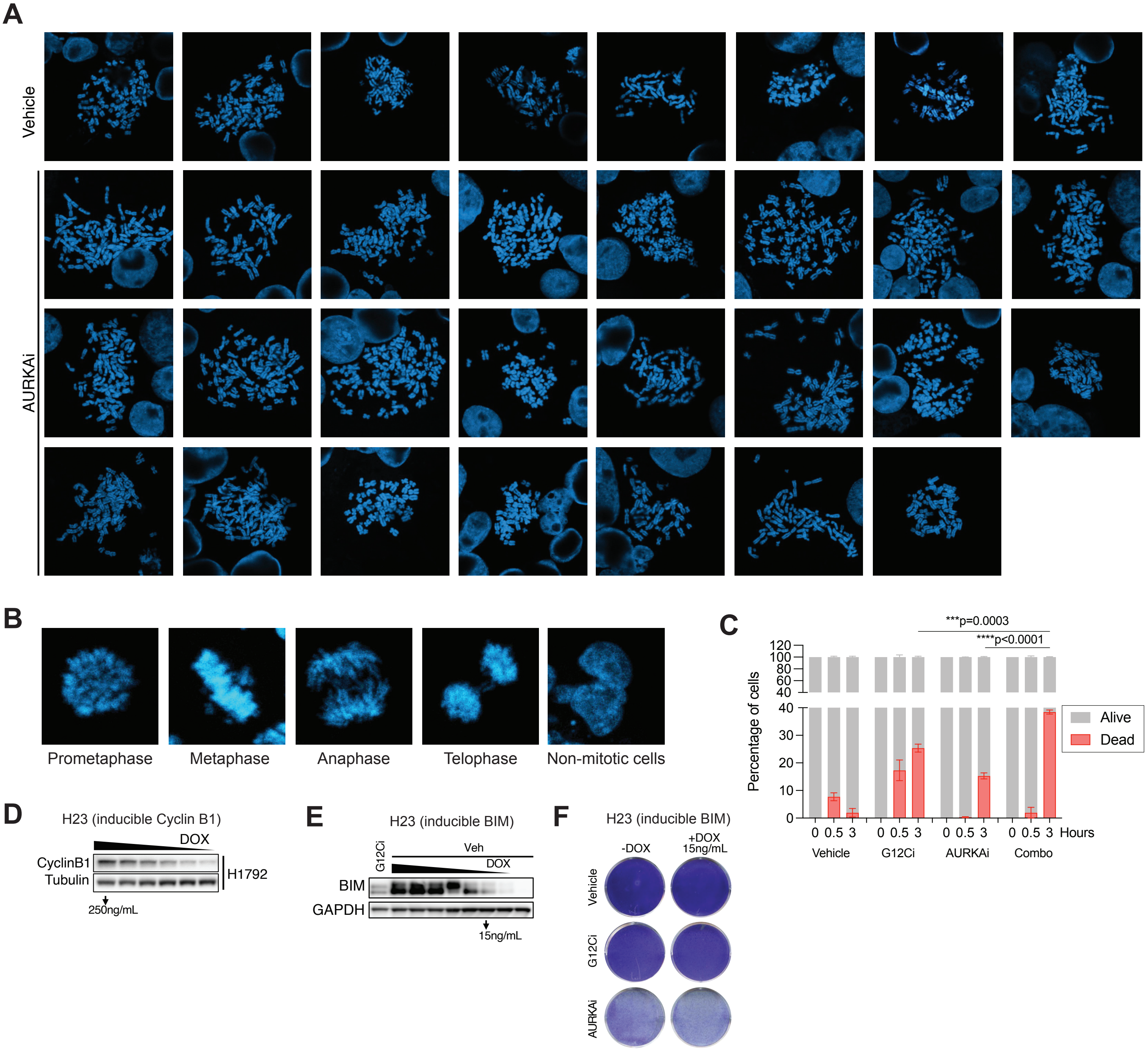


Supplemental Figure 5

### Supplemental Figure 6

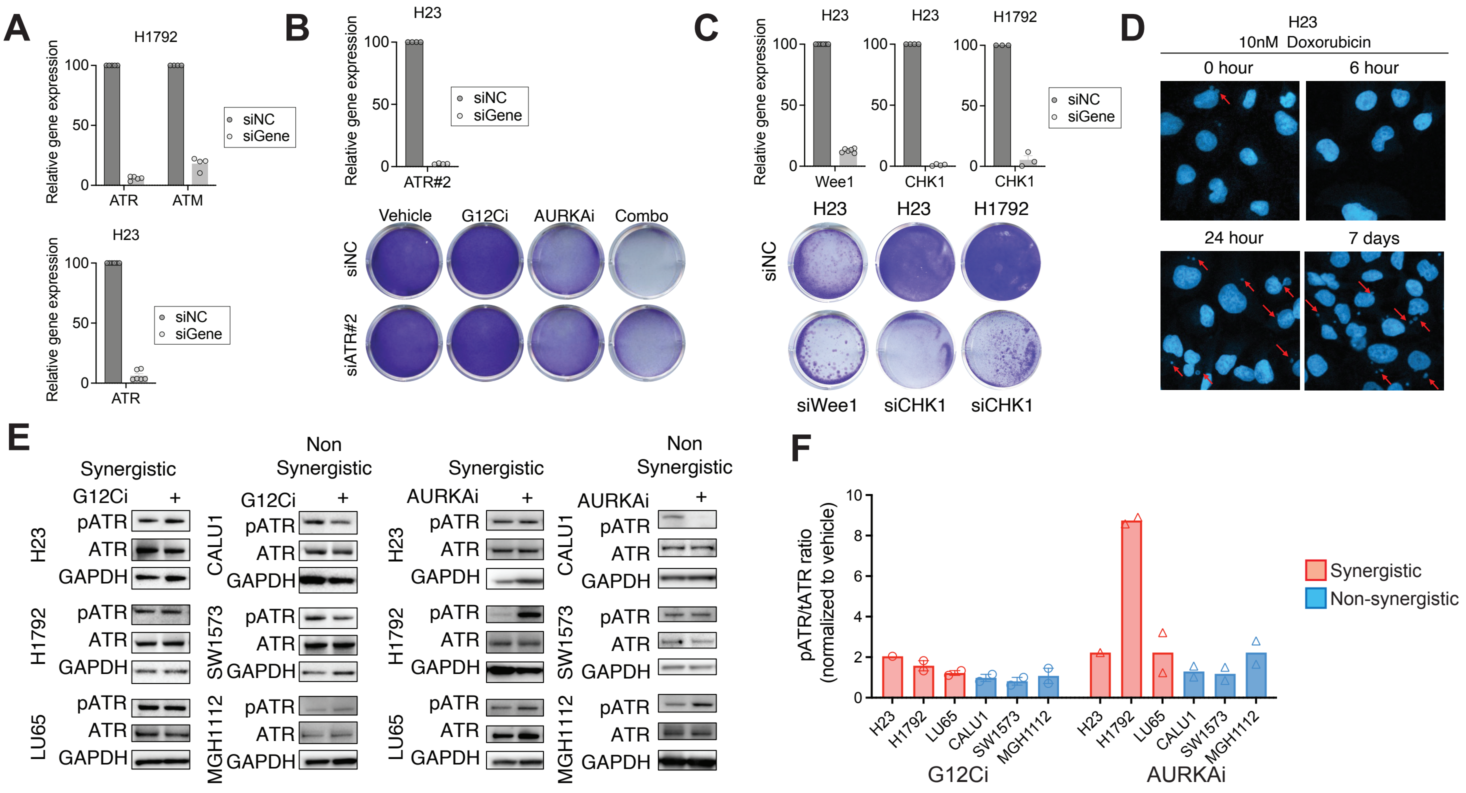


Supplemental Figure 6
